# Sparse autoencoder features from InterPLM predict neuropeptide precursors among secreted proteins

**DOI:** 10.64898/2026.08.20.746077

**Authors:** Anastasiya V. Kulikova, Angie L. Bookout, Thomas Lund Koch, Helena Safavi-Hemami

## Abstract

Neuropeptides are a diverse class of short, secreted signaling molecules that regulate key physiological processes in animals. Despite their important biological roles and increasingly recognized therapeutic value, the discovery of new neuropeptides remains challenging, largely because their short length and high sequence heterogeneity limit the effectiveness of motif- and homology-based approaches. Here, we present a pipeline for neuropeptide precursor prediction that leverages sparse autoencoders (SAEs) from InterPLM to decode dense ESM-2 protein language model embeddings into sparse, disentangled features. We identify a small subset of features strongly associated with neuropeptide precursors that achieve high discriminative performance. A logistic regression classifier trained on this reduced feature set, accurately separates human neuropeptide and non-neuropeptide sequences. We then applied this classifier to important model organisms: mouse (*Mus musculus*), zebrafish (*Danio rerio*), nematode (*Caenorhabditis elegans*), and fruit fly (*Drosophila melanogaster*) and show that the approach generalizes across diverse species. Overall, InterPLM SAE features provide an interpretable and effective strategy for neuropeptide prediction and enable a trained classifier to predict neuropeptides from large datasets. A web tool for this classifier is freely available at https://biolib.com/ATGCACTGTTCAGGCCTC/SAE-Neuropeptide-Predictor

## 1 Introduction

Neuropeptides and peptide hormones (hereafter collectively referred to as neuropeptides) are signaling molecules that regulate a wide range of physiological and behavioral processes, including synaptic transmission, growth, metabolism, learning, social behavior, and circadian rhythms [6, 20, 31]. Dysregulation of neuropeptide signaling is involved in numerous neurological and psychiatric disorders, such as chronic pain, depression, anxiety, neurodegenerative diseases, and metabolic syndromes [6]. Consequently, neuropeptides represent attractive therapeutic agents for various diseases [6].

While human neuropeptides have been extensively studied over the past decades, recent advances in gene prediction, proteomics, transcriptomics, and ribosome profiling suggest that many neuropeptides remain to be discovered in humans and other animals. Despite this progress, identifying novel neuropeptides remains challenging due to their distinct sequence and structural characteristics.

Neuropeptides are generated through proteolytic processing of precursor proteins and frequently undergo post-translational modifications, including disulfide bond formation, N-terminal glutamate cyclization, and C-terminal amidation [15]. Most neuropeptide precursors share a common organization with an N-terminal signal peptide that facilitates entry into the secretory pathway, followed by one or more propeptide regions that contain the neuropeptide sequence. Some precursors encode a single peptide, whereas others give rise to multiple peptides. For example, the precursors of insulin and relaxin each contain one neuropeptide, while the glucagon precursor is processed into several peptides, including glucagon, glucagon-like peptide 1, and oxyntomodulin.

Historically, neuropeptide discovery relied on biochemical isolation of mature neuropeptides from tissue extracts. In contrast, contemporary sequencing-based approaches typically begin with the identification of precursor genes or transcripts, followed by computational prediction and/or proteomic validation of the resulting peptides. A central challenge in this workflow is that neuropeptide precursors are often short, sequence-divergent, and lack well-defined structural domains. Consequently, approaches such as sequence homology-based searches, domain scans, or motif-based methods often fail to generalize beyond well-known families and miss more divergent sequences that lack recognizable motifs [20, 6]. Moreover, large-scale sequence-based discovery approaches often generate numerous candidates [5, 21], making it difficult to distinguish bona fide neuropeptides from other peptides and proteins or putative translated products that lack neuropeptide function. Because of these limitations, computational strategies that can learn and search for neuropeptide precursor-like properties without relying on fixed rules are promising for neuropeptide precursor prediction. Recent machine-learning approaches have begun to address this challenge by leveraging learned sequence representations (embeddings) to characterize neuropeptides and other proteins [9, 40, 13, 14, 43]. In particular, protein language models (LLMs) such as ESM-2 provide powerful protein embedding representations that capture biochemical and functional information directly from sequence [16, 26, 4], enabling the development of tools such as DeepPeptide for neuropeptide discovery [35]. However, a key limitation is that LLM embeddings are difficult to interpret because biological concepts can be distributed across many overlapping dimensions [4, 1, 37]. To address this challenge, InterPLM recently introduced sparse autoencoders (SAEs) trained on ESM-2 embeddings to disentangle overlapping signals into sparse, more interpretable features [29, 1, 28]. Notably, these features include signals associated with neuropeptide-like properties, providing a promising avenue for improving both the interpretability and performance of neuropeptide discovery approaches.

A key requirement for pursuing this avenue is a clear operational definition of what constitutes a neuropeptide. In this study, we define neuropeptides as peptides that are proteolytically processed from a secreted precursor protein beyond the initial removal of the N-terminal signal peptide. This definition encompasses most known neuropeptide diversity while excluding larger peptide hormones, such as growth hormone, prolactin, and erythropoietin, which typically do not undergo proteolytic processing following signal peptide removal. These hormones are also substantially larger than classical neuropeptides such as oxytocin or neurotensin and are therefore more readily identifiable using homology-based approaches.

Because neuropeptides are, by this definition, derived from precursor proteins, our analyses focus on identifying neuropeptide precursors that encode one or more mature neuropeptides. For simplicity, we hereafter refer to these precursor sequences as neuropeptides unless otherwise stated. Specifically, we leverage InterPLM SAE features to build a pipeline for neuropeptide precursor prediction and prioritization. We first embed secreted proteins (the secretome), including neuropeptide-encoding precursors, using ESM-2 [16] and encode these representations with an InterPLM SAE to obtain sparse feature activations [29]. We then quantify how strongly each SAE feature is associated with neuropeptide labels by scoring features across a set of thresholds and ranking them by their *F*_1_ scores. We then train a logistic regression classifier using these top-ranked features and evaluate the model on a curated human secretome dataset. We show that high-probability predictions are strongly enriched for annotated neuropeptides and that our classifier outperforms two existing neuropeptide prediction models [35, 11]. Finally, we test the approach on four model organisms, *M. musculus*, *D. rerio*, *C. elegans*, and *D. melanogaster*, demonstrating that interpretable SAE-based features support neuropeptide prediction across diverse species.

## 2 Methods

### 2.1 Training dataset collection

A positive and negative dataset were compiled to rank InterPLM features based on their association with neuropeptides, and this same dataset was later used to train the final classifiers.

To construct the positive training set for feature ranking and neuropeptide classification, sequences were combined from UniProtKB and OrthoDB [36, 32]. The initial dataset included 10,734 UniProtKB entries annotated with the neuropeptide keyword (KW-0527), 32,492 OrthoDB entries representing orthologs of G protein-coupled receptor (GPCR)-binding peptide families (accessed October 2024), and 9,994 OrthoDB ortholog sets corresponding to major peptide hormone families that do not target GPCRs (accessed October 2025), yielding 53,220 total sequences (*n* = 50,025 after deduplication).

To prevent train–test leakage, homologs to the human secretome test set were removed using MMseqs2 (*>* 80% sequence identity at *≥* 70% coverage), eliminating 15,316 sequences [30]. Next, DeepTMHMM-1.0 was applied to remove potential receptors or membrane proteins, removing an additional 921 sequences (*n* = 45,033) [7]. Finally, the remaining positives were partitioned with GraphPart (v1.0.2) into 10 folds at 40% identity to reduce overrepresentation of closely related peptides and enforce sequence diversity across partitions, resulting in a final positive training set of 35,483 sequences [34].

To construct the negative training set, all UniProtKB secreted proteins (KW-0964) were downloaded, yielding 2,168,324 sequences [36]. MMseqs2 was used to remove proteins homologous to the positive training set (*>* 40% sequence identity at *≥>* 70% coverage), removing 69,875 sequences and leaving 2,086,255 proteins [30]. Homologs to the human secretome test set were then removed with MMseqs2 (*>* 80% identity at *≥* 70% coverage), eliminating 75,384 additional sequences and leaving 2,010,871 proteins [30]. To obtain a diverse negative set, the remaining proteins were clustered with MMseqs2 at 40% identity and one representative sequence per cluster was retained, resulting in 102,288 sequences [30]. Finally, to match the length distribution of the positive dataset, both positive and negative sequences were binned by sequence length into 50-residue intervals, and negatives were randomly sampled within each bin (seed = 9,437,569) to match the positive bin counts, yielding a final negative training set of 33,615 sequences.

### 2.2 Test dataset collection

To construct a human test dataset enriched for secreted proteins while removing common non-neuropeptide classes, secreted human proteins were downloaded from UniProtKB using the query taxonomy id:9606 AND keyword:KW-0964, yielding 19,665 sequences [36]. The dataset was then deduplicated and filtered to remove transmembrane proteins, enzymes, and immunoglobulins, which are common in secreted-protein databases but do not constitute neuropeptides as per our definition.

To remove transmembrane proteins and enzymes, UniProtKB annotations were combined with homology-based filtering. First, transmembrane entries were identified using UniProtKB keyword and GO-term annotations (e.g., transmembrane keyword KW-0812 and membrane-associated GO terms) and added to a reference exclusion database unless they were annotated with hormone/neuropeptide activity GO terms (protected set). A reference set of proteins annotated with enzyme-related GO molecular function terms was also downloaded from UniProtKB and appended to the exclusion database. The secretome FASTA file was then searched against this exclusion set using MMseqs2, and sequences were removed if an MMseqs2 alignment satisfied: E-value *≤* 10*^−^*^10^, percent identity *≥* 35%, target coverage *≥* 60%, and aligned length *≥* 20 amino acids [30]. 15,316 sequences remained after transmembrane proteins and enzymes were removed (see the github repository for the full list of GO terms used).

Immunoglobulins were removed using a combination of UniProtKB annotation filters and homology-based screening. Secretome entries annotated with immunoglobulin-related terms (`GO:0019814; KW-0377`) were flagged, and a reference immunoglobulin query set was downloaded from UniProtKB using GO:0019814 and/or the immunoglobulin-domain keyword `KW-0377`. Sequences were removed if they matched the immunoglobulin reference database under the same MMseqs2 criteria used above [30]. After this filtering step, 1211 sequences remained.

To remove N-terminally truncated sequences and those annotated as secreted but lacking a signal peptide, only sequences with a SignalP 6.0 score *≥* 0.4 were retained [33]. Remaining sequences were clustered with CD-HIT at 95% identity to reduce redundancy and bias in test scoring [10]. After all filtering steps, the final human test dataset contained 562 sequences, which were then manually evaluated to yield 115 annotated neuropeptides. Manual validation included removing truncated sequences and excluding large peptide hormones that are not proteolytically cleaved from precursor proteins (e.g., growth hormone, prolactin). Predictions for the human secretome are provided in Supplementary Data File S1.

### 2.3 Collecting additional secretomes

Additional secretomes from four model organisms were collected from UniProtKB by querying for secreted proteins using organism-specific taxonomy identifiers: *D. rerio* (taxid 7955), *M. musculus* (taxid 10090), *C. elegans* (taxid 6239), and *D. melanogaster* (taxid 7227) [36]. The same general filtering workflow was applied across species, including deduplication, removal of transmembrane proteins, likely enzymes and (where applicable) immunoglobulins, SignalP filtering, CD-HIT clustering, and manual evaluation of annotations.

For the *M. musculus* secretome, the initial download contained 4,429 sequences. After deduplication and removal of enzymes and immunoglobulins, 948 sequences remained. SignalP filtering (score *≥* 0.4) reduced the set to 753 sequences, and CD-HIT clustering yielded a final set of 515 sequences. The final mouse dataset contained 125 manually annotated neuropeptides. Predictions for the *M. musculus* secretome are provided in Supplementary Data File S2.

For the *D. melanogaster* secretome, the initial dataset contained 1,361 sequences. After deduplication and removal of enzymes, 566 sequences remained. SignalP filtering reduced this to 464 sequences, and CD-HIT clustering yielded a final set of 265 sequences, including 28 annotated neuropeptides. Predictions for the *D. melanogaster* secretome are provided in Supplementary Data File S3.

For the *C. elegans* secretome, the initial download contained 477 sequences. After deduplication and removal of enzymes, 217 sequences remained, and SignalP filtering reduced the set to 197 sequences. Following CD-HIT clustering, 189 sequences remained; one unusually large protein (*>* 6,000 residues) was removed. The final worm dataset contained 36 annotated neuropeptides. Predictions for the *C. elegans* secretome are provided in Supplementary Data File S4.

For the *D. rerio* secretome, the initial download contained 1,945 sequences. After removing enzymes, 680 sequences remained, and deduplication yielded 663 sequences; no immunoglobulin filtering was applied. SignalP filtering (score *≥* 0.4) left 554 sequences. After CD-HIT clustering, the final dataset contained 454 sequences; one protein *>* 6,000 residues was removed, and 88 annotated neuropeptides were retained. Predictions for the *D. rerio* secretome are provided in Supplementary Data File S5.

### 2.4 Embedding generation

SAE feature embeddings were generated using the InterPLM sparse autoencoder, which produces sparse feature activations from ESM-2 sequence representations [29, 16]. For each protein sequence in the input FASTA file, residue-level embeddings were computed with the ESM-2 650M model (`esm2_t33_650M_UR50D`), and activations from layer 18 were extracted. These layer-18 embeddings were then passed through a pretrained sparse autoencoder for the corresponding ESM-2 model and layer (`plm_model=esm2-650m, plm_layer=18`), producing a matrix of per-residue SAE activations for each sequence (sequence length *×* feature dimension).

### 2.5 Calculating F_1_ scores

To quantify the association between individual InterPLM/SAE features and neuropeptide labels, perfeature *F*_1_ scores were calculated from max-pooled sequence embeddings. For each feature, per-residue activations were reduced to a 1-D vector by taking the maximum activation across the sequence. A set of thresholds (0.0, 0.15, 0.3, 0.4, 0.5, 0.6, and 0.8) was evaluated; at each threshold, a protein was called “positive” for that feature if its max activation exceeded the threshold. For every feature–threshold pair, we counted true positives (*TP*), false positives (*FP*), and false negatives (*FN*), computed precision and recall, and calculated

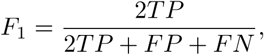

with *F*_1_ set to 0 when both precision and recall were 0. Finally, scores were max-pooled across thresholds by retaining the highest *F*_1_ score per feature and recording the corresponding threshold, and features were ranked by their maximum *F*_1_ score.

### 2.6 Classifier training and model selection

A logistic regression classifier was trained to predict neuropeptide versus non-neuropeptide sequences from max-pooled SAE feature embeddings. The classifier was trained on the same positive and negative datasets used for per-feature *F*_1_ score calculations. The positive dataset was split into 10 folds using GraphPart clusters (40% identity) to reduce homology-driven overrepresentation within splits [34]. Because the negative dataset was already clustered for diversity, it was divided evenly into 10 folds. Model selection was performed using nested 10-fold cross validation. In each outer iteration, one fold was held out as a test fold and the remaining nine folds were used for training and validation; feature rankings were computed using the training data only, preventing information leakage from the held-out fold.

Within each outer fold, an inner cross-validation loop was used to select how many top-ranked SAE features to include. Models were trained using increasing numbers of features (in steps of five, up to 1,000). For each feature count, models were trained on eight folds and validated on the ninth fold, repeating so each fold served as validation once; validation accuracies were averaged. Features were standardized to zero mean and unit variance, and logistic regression was fit with the L-BFGS solver (`max_iter=3000`; seed 9,687,254). Outer-fold test accuracy was obtained by retraining on all nine non-held-out folds and evaluating on the held-out test fold. Test accuracies were averaged across outer folds for each feature count, and the final number of features (320) was chosen using the elbow method from the Kneedle algorithm on the mean test-accuracy curve [27].

To train the final model, features were selected using a global ranking generated by combining all 10 folds, and a final logistic regression classifier was trained on the full training dataset using the top 320 features for downstream evaluation.

### 2.7 Secretome evaluation

To evaluate the final classifier on the human secretome test set, SAE feature embeddings were generated as described above and reduced to a single 1-D feature vector per protein by max pooling across residues. The final logistic regression model was applied using the top 320 features from the global feature ranking, producing a predicted neuropeptide probability for each sequence (0–1). A probability threshold of 0.7 was used to convert probabilities into binary predictions, and a confusion matrix was computed (TP, FP, TN, FN). Accuracy, precision, recall, and *F*_1_ score were calculated from these counts. The same evaluation procedure was applied to the *M. musculus*, *D. melanogaster*, *C. elegans*, and *D. rerio* secretomes. Predictions for all species are provided in Supplementary Data File S1, Supplementary Data File S2, Supplementary Data File S3, Supplementary Data File S4, and Supplementary Data File S5.

### 2.8 Principal component analysis

To assess whether neuropeptides occupy distinct regions of embedding space under unsupervised dimensionality reduction, we performed principal component analysis (PCA) on human secretome protein embeddings derived from ESM-2 and InterPLM (SAE) representations under two pooling strategies (mean and max). For each embedding type, PCA was computed and proteins were projected into the first two principal components (PC1 and PC2). Points were labeled as “Neuropeptide” or “Non-neuropeptide” using the curated annotations, and separation between the two groups in PC1–PC2 space was quantified using centroid distance and the silhouette score.

Centroid separation was computed by first finding the average PC1 and PC2 values for neuropeptides and for non-neuropeptides (i.e., the “center” of each group in the PC1–PC2 plot), and then measuring the Euclidean distance between these two centers. If the neuropeptide centroid is (*PC*1*_NP_, PC*2*_NP_*) and the non-neuropeptide centroid is (*PC*1*_Non_, PC*2*_Non_*), the centroid distance was calculated as

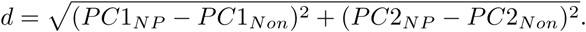

To complement centroid separation with a neighborhood-based measure of cluster separation, we computed the mean silhouette score in PC1–PC2 space. Pairwise Euclidean distances were calculated on the PC1–PC2 coordinates, class labels were encoded as integers, and silhouette widths were computed using the cluster package in R. The mean silhouette score across all points was reported for each embedding condition, with values near 0 indicating substantial overlap between classes and negative values indicating that points are, on average, closer to the opposite class than to their assigned class.

### 2.9 Benchmarking against other models

To benchmark our InterPLM (SAE) logistic regression classifier against existing neuropeptide prediction tools, we evaluated two published methods on the curated human secretome test set, DeepPeptide and NeuroPP [35, 11]. DeepPeptide predictions were obtained by running the model and parsing its JSON output; we then converted these outputs into a binary call by labeling a precursor as positive if at least one mature peptide was predicted. NeuroPP predictions were obtained from the NeuroPP web server and evaluated using a probability threshold of *≥* 0.5 to generate binary labels. For each baseline model, the resulting binary predictions were compared to our human secretome predictions thresholded at *≥* 0.7 to compute confusion matrices and summary metrics (accuracy, precision, recall, and *F*_1_ score). A threshold of *≥* 0.7 was chosen for our model as most correctly predicted neuropeptides have probabilities above 0.7 as shown in Fig. 5a. As a supplementary analysis, two additional models were tested, NeuroPred-FRL and NeuroPred-PLM [9, 40]. NeuroPred-FRL predictions were obtained by submitting human secretome sequences to the NeuroPred-FRL online web server, which returns a binary classification for each sequence (1 indicating neuropeptide; 0 indicating non-neuropeptide) [9]. NeuroPredPLM predictions were generated locally by installing the NeuroPred-PLM software package (v0.1.0) and running it on the secretome sequences. This model also outputs a binary prediction for each sequence (1/0) [40].

### 2.10 Data availability

Final data analysis and figure production were performed in R [25], using the tidyverse family of packages [42]. Data processing and modeling were performed in Python using NumPy [8], pandas [18], PyTorch [23], scikit-learn [24], Biopython [3], and the InterPLM framework [29]. Analysis scripts and processed data are available on GitHub: https://github.com/akulikova64/neuropeptide_SAE_project.

The trained classifier is available as a Python package and can be installed locally from PyPI: https://pypi.org/project/sae-neuropeptide-predictor/. Source code for the predictor is available at https://github.com/akulikova64/sae-neuropeptide-predictor. A web-based tool for the classifier is also available through BioLib: https://biolib.com/ATGCACTGTTCAGGCCTC/SAE-Neuropeptide-Predictor.

## 3 Results

### 3.1 Neuropeptide precursors show enrichment in InterPLM PCA space

To compare whether dense ESM-2 embeddings or InterPLM (SAE) embeddings better capture neuropeptide-related structure, we performed principal component analysis (PCA) on human secretome protein embeddings using both mean-pooled and max-pooled embeddings across residues (Fig. 1a–d). Across all conditions, proteins annotated with neuropeptide labels and other secreted proteins substantially overlapped in PC1–PC2 space, indicating that neuropeptides are not cleanly separable in a two-dimensional projection. However, the InterPLM projections had a pronounced triangular shape and neuropeptides were enriched near an extreme vertex (the “tip” of the triangle), suggesting that neuropeptides are shifted toward one edge of the InterPLM representation space rather than forming a distinct isolated cluster. This shift indicates a neuropeptide-associated signal in the embedding space that may be weak in a 2D projection but may still be leveraged for discrimination when higher-dimensional features are used together.

**Figure 1:**
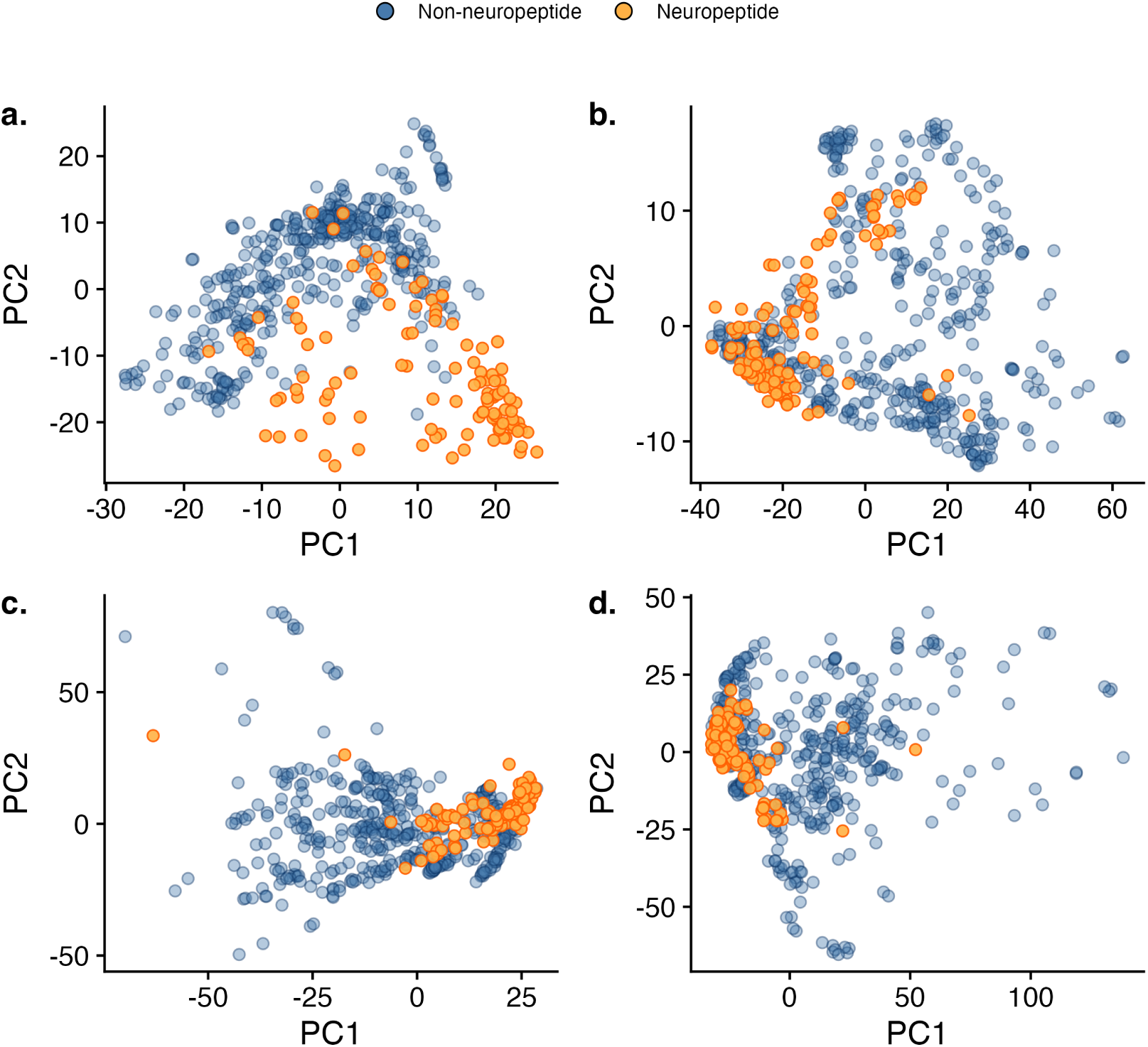
Principal component analysis (PCA) of secretome protein embeddings for the ESM and InterPLM models with different pooling strategies. PCA projections are shown for (a) averaged ESM-2 embeddings, (b) max-pooled ESM-2 embeddings, (c) averaged InterPLM embeddings, and (d) max-pooled InterPLM embeddings. Orange points indicate annotated neuropeptides, while blue points indicate non-neuropeptide secreted peptides. Centroid distances between neuropeptides and other secreted peptides in PC1–PC2 space were 21.01 for averaged ESM-2 embeddings and 23.08 for averaged SAE embeddings. Max-pooled ESM-2 embeddings showed a centroid distance of 20.94, while max-pooled SAE embeddings showed a centroid distance of 28.38. Silhouette scores in PC1–PC2 space were small across all conditions (ESM-2 averaged: 0.35; ESM-2 maxpooled: 0.06; SAE averaged: 0.07; SAE maxpooled: *−*0.01).

InterPLM embeddings showed larger centroid distances between neuropeptides and other secreted proteins than ESM-2, particularly under max pooling (Fig. 1). Centroid distances in PC1–PC2 space were 21.01 (ESM-2 mean) and 20.94 (ESM-2 max) compared to 23.08 (InterPLM mean) and 28.38

(InterPLM max) (Fig. 1). Silhouette scores remained small across conditions (ESM-2 mean: 0.35; ESM-2 max: 0.06; InterPLM mean: 0.07; InterPLM max: *−*0.01), consistent with substantial overlap despite the larger centroid distances.

Although neuropeptides are not cleanly separated in a 2D PCA projection, their enrichment near an extreme region of the InterPLM embedding space and the increased centroid separation suggest that the representation contains predictive signal, supporting the training of a classifier for improved class separation. Throughout the rest of the paper, we used max-pooled L18 InterPLM (SAE) embeddings.

### 3.2 Top InterPLM features show strong association with neuropeptides

To identify InterPLM features most associated with neuropeptide labels (Fig. 2), we curated positive and negative training sets designed to be diverse and non-overlapping with the human secretome as a test set (Fig. 2a,b). We then generated layer 18 InterPLM embeddings for each sequence and max-pooled across residues to obtain a single 10,240-dimensional feature vector per protein (Fig. 2c). For every feature, we evaluated a simple binary classifier across seven candidate activation thresholds (0, 0.15, 0.3, 0.4, 0.5, 0.6, and 0.8), scoring how well that feature alone recovered the positive set while avoiding false positives in the negative set (*F*_1_ score) (Fig. 2d). Finally, we kept the best *F*_1_ score (and its corresponding threshold) for each feature and ranked features by this maximum to prioritize neuropeptide-associated features for downstream classifier training (Fig. 2e).

**Figure 2:**
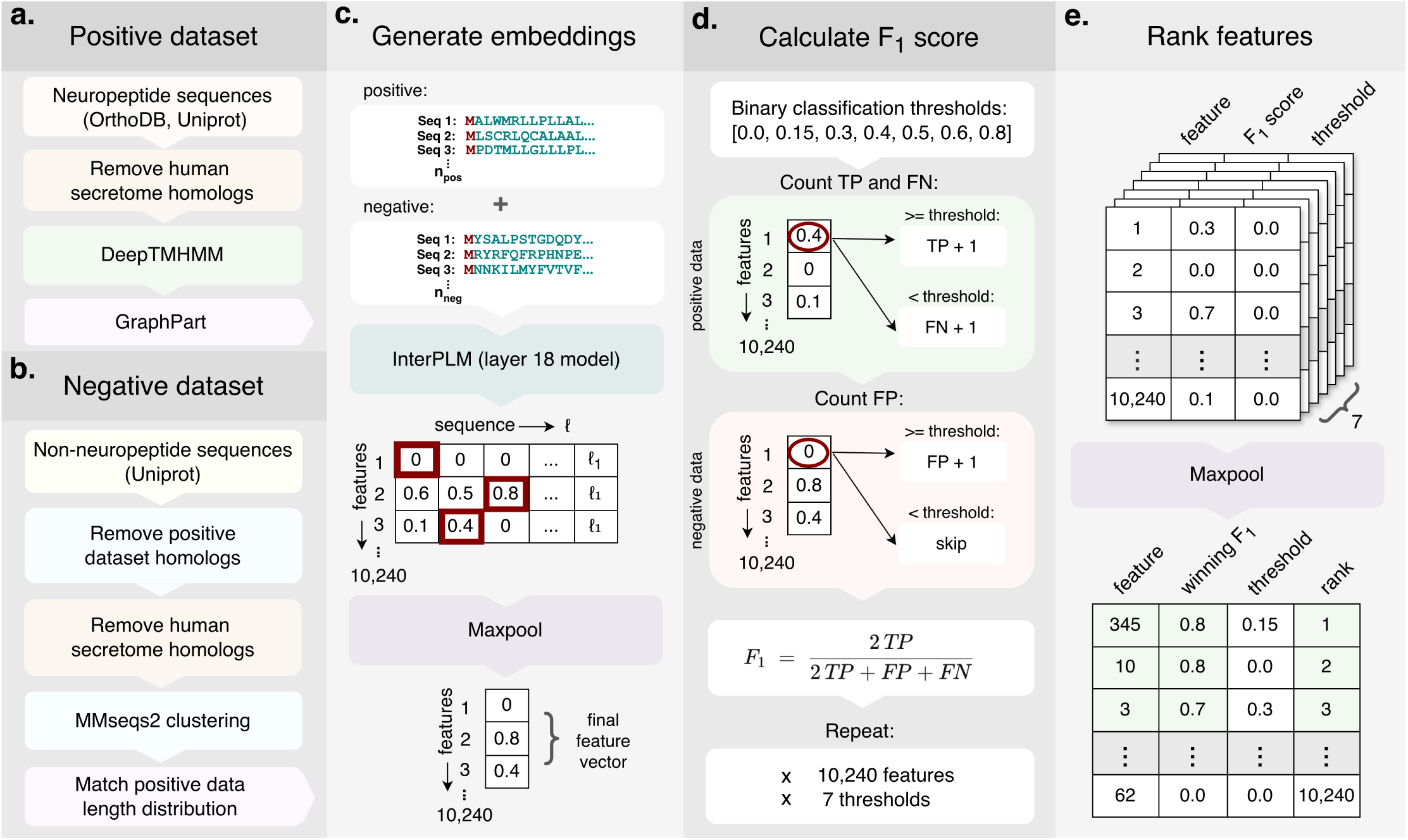
Pipeline for identifying neuropeptide-associated features across layer 18 InterPLM embeddings. (a) Pipline for curating positive training data. (b) Pipline for curating negative training data. (c) Embedding generation. Embeddings are generated for both the positive and negative datasets. Layer 18 embeddings are maxpooled across the sequence resulting in a 1-D vector of 10,240 features. (d) Calculating *F*_1_ score. Binary classifier uses 7 thresholds to calculate *F*_1_ score across all features using the positive (green) and negative (pink) maxpooled embeddings. (e) Feature ranking. Features are maxpooled across 7 thresholds and ranked based on decending *F*_1_ score (feature with highest *F*_1_ is ranked as the top feature)

Across max-pooled embeddings, InterPLM features demonstrate significantly stronger and more informative associations with neuropeptides than raw ESM-2 features (Fig. 3). The distribution of per-feature *F*_1_ scores for ESM-2 demonstrates degeneracy, with many features collapsing to a similar value (*∼* 0.66), whereas InterPLM (SAE) produces a broad distribution with many features achieving high *F*_1_ scores (often *>* 0.8; Fig. 3a). Consistent with this difference, the thresholds at which features achieved their best *F*_1_ score were largely concentrated at the highest tested threshold for ESM-2 (0.8), while InterPLM (SAE) features more frequently peaked at lower thresholds (Fig. 3b), suggesting that neuropeptide-relevant signals are captured as stronger activations in the sparse feature space. InterPLM (SAE) also yielded substantially more features with *F*_1_ *≥* 0.5 than ESM-2 (Fig. 3c); however, many of the ESM-2 features with *F*_1_ *∼* 0.66 reflect degenerate behavior at low activation thresholds. Finally, inspection of the top-ranked InterPLM (SAE) features highlighted a small subset of highly predictive features and their associated optimal thresholds (Fig. 3d). Most of these high-performing features achieved their optimal *F*_1_ score at a threshold of zero, indicating stronger, less noisy discrimination between neuropeptides and non-neuropeptides. Importantly, none of the top-ranked features required high thresholds to achieve strong *F*_1_ performance, in contrast to degenerate scenarios (such as with ESM-2 features) in which apparent performance can arise from predicting only a minimal number of high-confidence positives. Together, these results indicate that top-ranked InterPLM features show strong predictive performance individually, and combining them as inputs to a classifier should further improve class separation and overall prediction accuracy.

**Figure 3:**
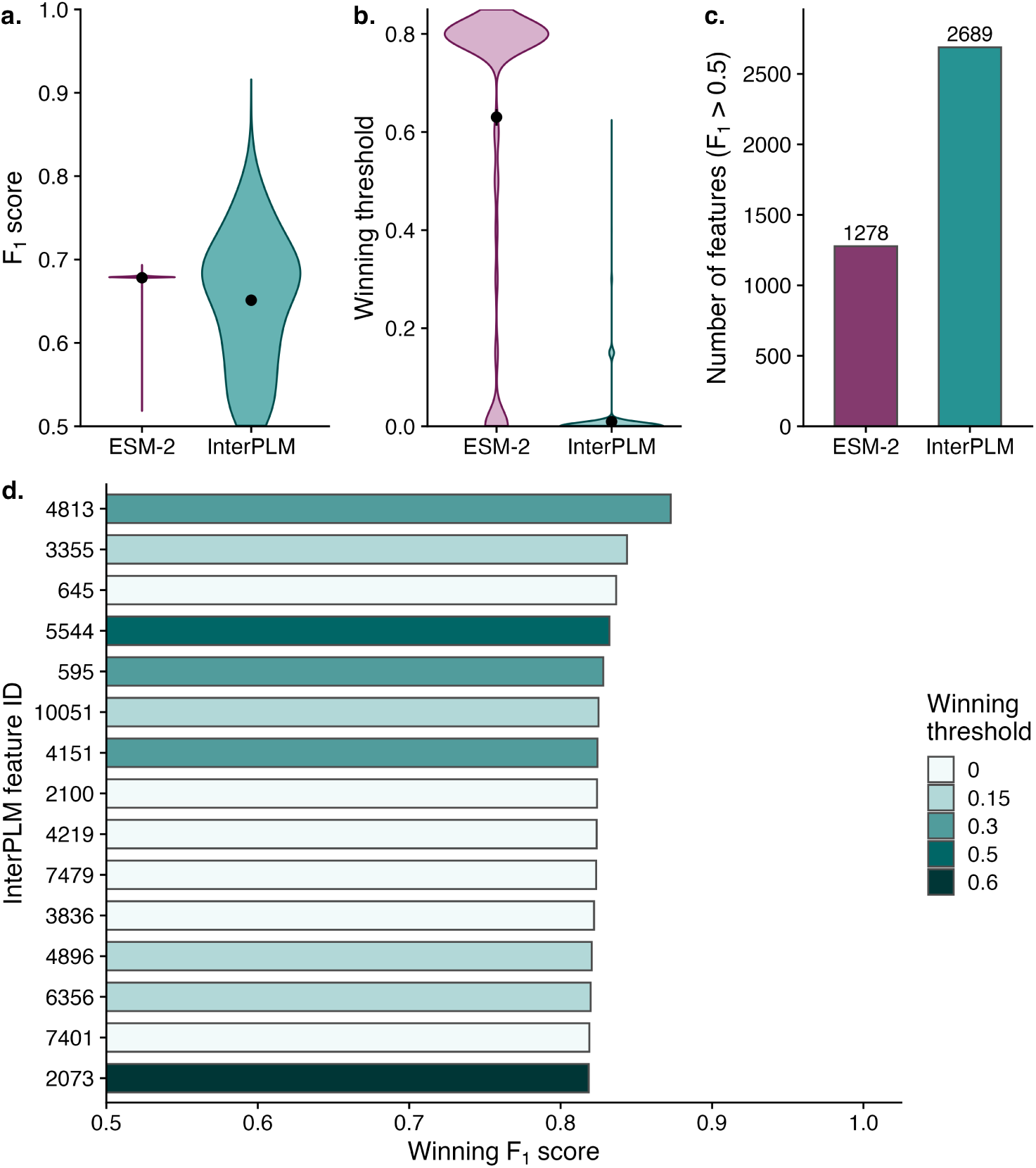
Per-feature *F*_1_ score analysis for max-pooled ESM-2 and InterPLM (SAE) embeddings. (a) Distributions of per-feature *F*_1_ scores. (b) Distributions of the thresholds yielding the maximum *F*_1_ score for each feature. (c) The number of features with *F*_1_ *≥* 0.5 per model. (d) The top 15 InterPLM (SAE) features ranked by maximum *F*_1_ score along with their corresponding optimal thresholds.

### 3.3 Logistic regression classifier identifies human neuropeptides

We next trained a logistic regression classifier in the SAE feature space and selected the final model size using nested 10-fold cross validation (Fig. 4). Positive sequences were assigned to 10 folds using GraphPart clustering (40% identity) to reduce homology-driven overrepresentation and to minimize data leakage from closely related sequences appearing in both training and test folds, whereas negatives were split evenly into 10 folds (Fig. 4a). For each outer test fold, feature rankings were recomputed from the remaining nine folds, yielding fold-specific rankings across all train/validation combinations (10 ranking files total; Fig. 4a,d). Logistic regression models were then trained on standardized, max-pooled SAE feature vectors using increasing numbers of top-ranked features in increments of 5 up to 1000 (Fig. 4b), and test performance was averaged across outer folds for each feature count *N* to produce an average test-accuracy curve (Supplementary Figs. S1 and S2). The final model size of 320 features was chosen as the number of features at the elbow of this curve using the Kneedle algorithm (Fig. 4c). Finally, all folds were combined to generate a global feature ranking and a final model was trained on the top *N* features (Fig. 4d,e). A PCA using only these top 320 neuropeptide-associated SAE features is shown in Supplementary Fig. S3. The PCA showed partial class separation where annotated neuropeptides occupy a more compact region of the top-320-feature space. However, substantial overlap with non-neuropeptides still remained.

**Figure 4:**
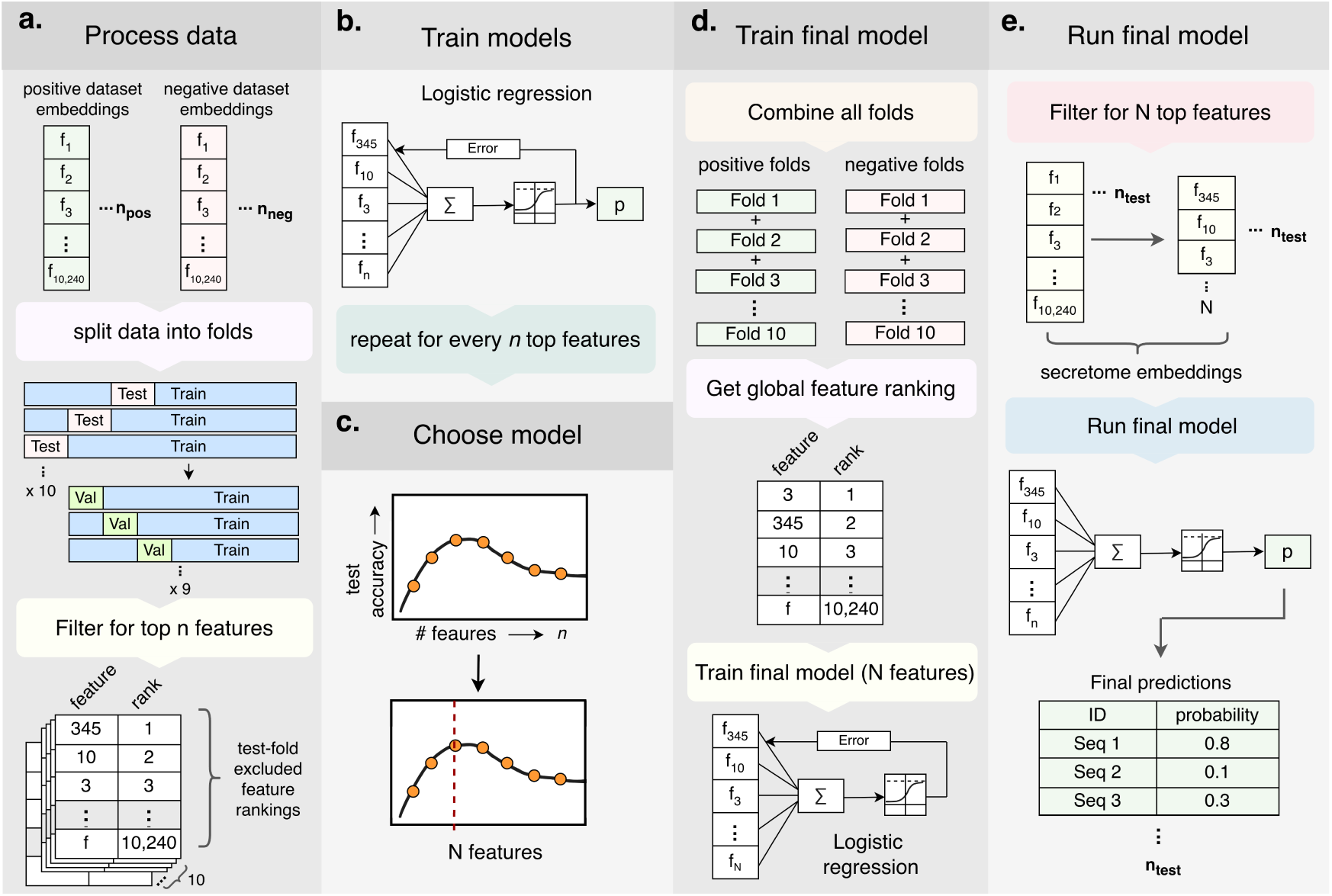
Classifier training, model selection, and secretome evaluation workflow. (a) Data preprocessing and partitioning. Positive sequences were assigned to 10 folds using GraphPart clustering, negatives were split into 10 folds evenly, and nested 10-fold cross validation was performed. Feature rankings (10 ranking files) were generated from all combinations of 9 validation folds excluding the test fold. (b) Logistic regression training. Max-pooled SAE feature vectors were used to train logistic regression models across increasing numbers of top-ranked features. (c) Model selection. Test performance was averaged across test folds for each feature count *N*, and the final model size was chosen from the average test-accuracy curve (kneedle elbow). (d) Training the final model. All folds were combined. Feature rankings were generated from all 10 folds to form a global ranking. Model was trained on the top N features from the global ranking. (e) Running the final model. The final model was applied to the secretome test dataset to output per-sequence neuropeptide probabilities.

We next evaluated the final InterPLM (SAE) logistic regression model, referred to as InterPLM-NP (Neuropeptide Predictor) throughout the rest of the paper, on the human secretome and examined how well the model can predict our manually annotated set of neuropeptides within the human secretome (Fig. 5). When predictions were binned into 10% probability intervals, the fraction of annotated neuropeptides increased sharply with increasing predicted probability, with the 90–100% bin containing 84% neuropeptides (Fig. 5a). This enrichment indicates that the model’s probability output provides a useful ranking of candidates, where high-scoring sequences are more likely to be true neuropeptides. Similarly, most annotated neuropeptides were concentrated in the highest-probability bin: of the 115 annotated neuropeptides in the human secretome, 90 received predicted probabilities above 0.9 (Fig. 5b). The remaining annotated neuropeptides were distributed across lower-probability bins. Consistent with this high-confidence enrichment, several well-characterized human peptide hormones were assigned predicted probabilities near 1.0 (Table 2).

**Figure 5:**
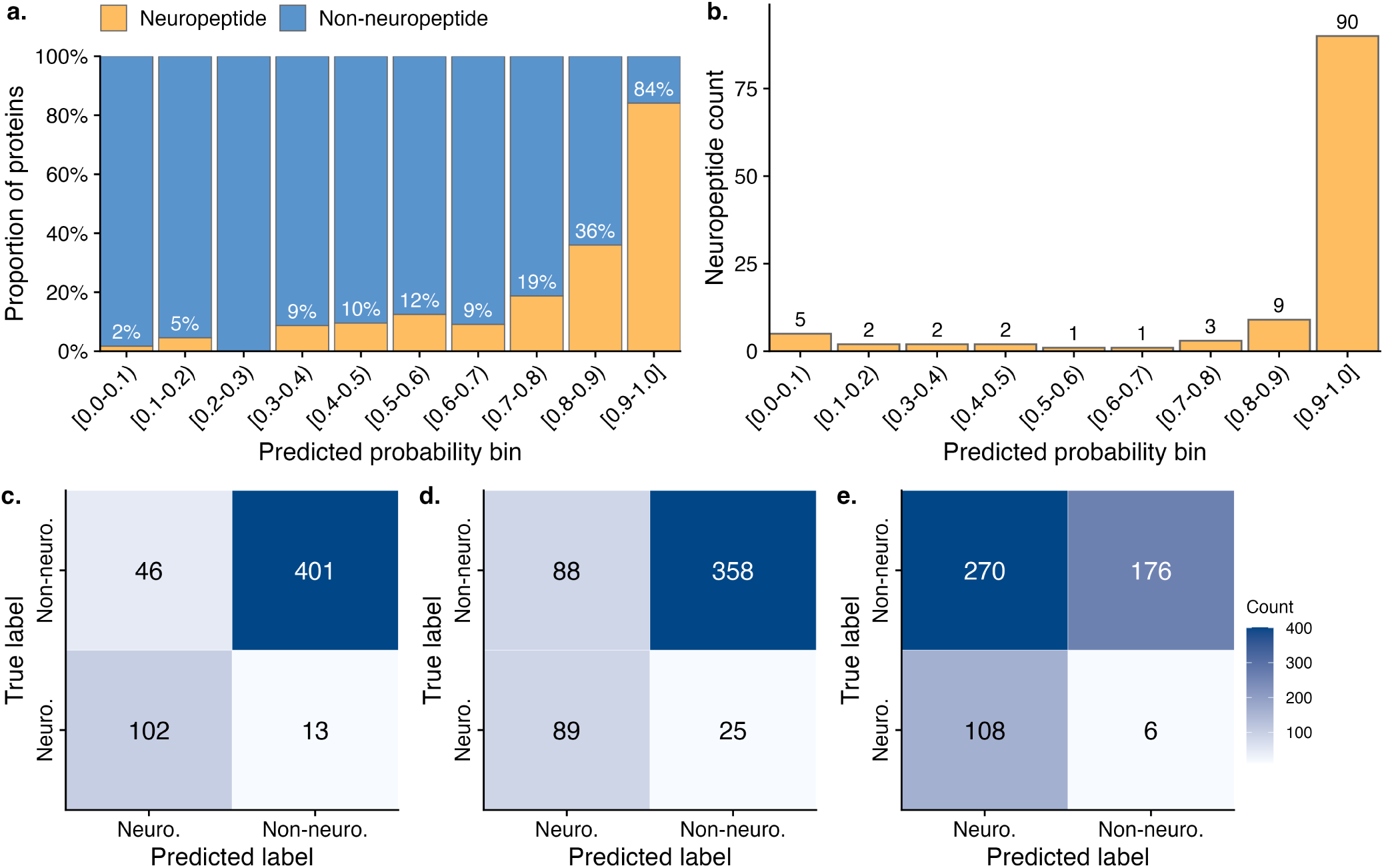
Classifier assessment and model comparison on the human secretome. (a) Proportion of annotated neuropeptides within 10% probability bins of the InterPLM-NP model predictions. (b) Counts of annotated neuropeptides across the probability bins. (c) Confusion matrix for InterPLM-NP predictions evaluated at a probability threshold of *≥* 0.7. (d) Confusion matrix for DeepPeptide predictions, where a precursor was called positive if at least one mature peptide was predicted. (e) Confusion matrix for NeuroPP predictions evaluated at a probability threshold of *≥* 0.5.

It is important to note that the true test accuracy assessment in this study comes from the nested cross-validation procedure used during model selection, where each outer fold served as a held-out test set and performance was averaged across folds (Supplementary Fig. S2). Because folds were made to reduce homology overlap (GraphPart clustering for positives and diversity clustering for negatives), the resulting test accuracies provide the closest approximation of true out-of-sample performance under controlled splits. By contrast, the human secretome evaluation should be viewed primarily as a realistic application scenario. Although we removed close homologs to the secretome from training at high identity (*>* 80%), secretome proteins may still share weaker similarity to training sequences and therefore are not guaranteed to be fully independent. As a result, the secretome analysis should be interpreted as a demonstration of how the classifier prioritizes candidates in a real-world secreted-protein set, whereas the cross-validation test curve provides the primary estimate of expected predictive performance which peaks at an average test accuracy of *∼* 92% at 320 features (Supplementary Fig. S2).

Finally, we compared the InterPLM-NP against two additional neuropeptide discovery tools, DeepPeptide and NeuroPP (Fig. 5c–e and Table 1). Using a probability threshold of *≥* 0.7, the InterPLM-NP recovered *∼* 88.7% annotated neuropeptides (TP=102; FN=13) while maintaining a moderate false-positive count (FP=46; TN=401; Fig. 5c). Since DeepPeptide predicts mature peptides within the precursors, we called a precursor as positive (neuropeptide) if at least one mature peptide was predicted. DeepPeptide identified *∼* 78.0% of annotated neuropeptides (TP=89; FN=25), but with more false positives (FP=88; TN=358; Fig. 5d). NeuroPP achieved very high recall (TP=108; FN=6) but produced substantially more false positives compared to the other two models (FP=270; TN=176; Fig. 5e) when evaluated at a probability threshold of *≥* 0.5. To assess agreement among the three models, we compared the sets of annotated neuropeptides each model correctly recovered. We found that 82 of the 115 annotated neuropeptides were identified by all three models, while only two were missed by all three (Supplementary Fig. S6). Additional baseline comparisons are provided in Supplementary Fig. S5. NeuroPP scores showed a moderate correlation with InterPLM-NP (*r* = 0.447, *p* = 6.96*×*10*^−^*^29^), showing strong agreement for a subset of high-confidence neuropeptides but substantial disagreement for many low-scoring InterPLM-NP sequences. We also report confusion matrices for NeuroPred-FRL and NeuroPred-PLM on the same secretome (Supplementary Fig. S5b,c and Table 1). However, these tools were trained for mature neuropeptide prediction rather than precursor-level classification, so their outputs are included primarily to contextualize how mature-peptide predictors behave on a precursor-labeled benchmark.

**Table 1:** Performance comparison of neuropeptide prediction models on the human secretome.

| Model | $F_1$-score | Accuracy | Precision | Recall |
| --- | --- | --- | --- | --- |
| **L18 InterPLM LogReg** | **0.776** | **0.895** | **0.689** | **0.887** |
| DeepPeptide | 0.612 | 0.798 | 0.503 | 0.781 |
| NeuroPP | 0.439 | 0.507 | 0.286 | 0.947 |
| NeuropredPLM | 0.233 | 0.673 | 0.224 | 0.243 |
| NeuropredFRL | 0.091 | 0.751 | 0.179 | 0.061 |

**Table 2:** Predicted neuropeptide probabilities for selected human peptide hormones.

| Accession | Probability | Neuropeptide |
| --- | --- | --- |
| A6XND7 | 0.999 | Pro-opiomelanocortin |
| P22466 | 0.998 | Galanin |
| P06307 | 0.998 | Cholecystokinin |
| P61278 | 0.998 | Somatostatin |
| P01282 | 0.997 | Vasoactive intestinal peptide |
| P01178 | 0.996 | Oxytocin-neurophysin 1 |
| O15130 | 0.996 | Neuropeptide FF |
| P01303 | 0.995 | Pro-neuropeptide Y |
| A0A0S2Z478 | 0.991 | Corticoliberin |
| G3V1X6 | 0.990 | Neurotensin/neuromedin N |
| P0C0P6 | 0.989 | Neuropeptide S |
| P01258 | 0.985 | Calcitonin |
| P01344 | 0.984 | Insulin-like growth factor 2 |
| P01350 | 0.981 | Gastrin |
| A6XGL2 | 0.975 | Insulin |
| P01185 | 0.968 | Vasopressin |
| P05019 | 0.956 | Insulin-like growth factor 1 |

### 3.4 Classifier identifies neuropeptides in other species

To assess cross-species generalization, we applied InterPLM-NP to four additional secretomes from *M. musculus*, *D. rerio*, *C. elegans*, and *D. melanogaster* (Fig. 6). Predicted probability distributions showed clear separation between annotated neuropeptides and non-neuropeptide secreted proteins in all four species, with non-neuropeptides concentrated near zero probability and neuropeptides concentrated above 0.9 probability (Fig. 6a). In *C. elegans* and *D. melanogaster*, annotated neuropeptides were predominantly high scoring, producing unimodal neuropeptide distributions that were well separated from the background secretome.

**Figure 6:**
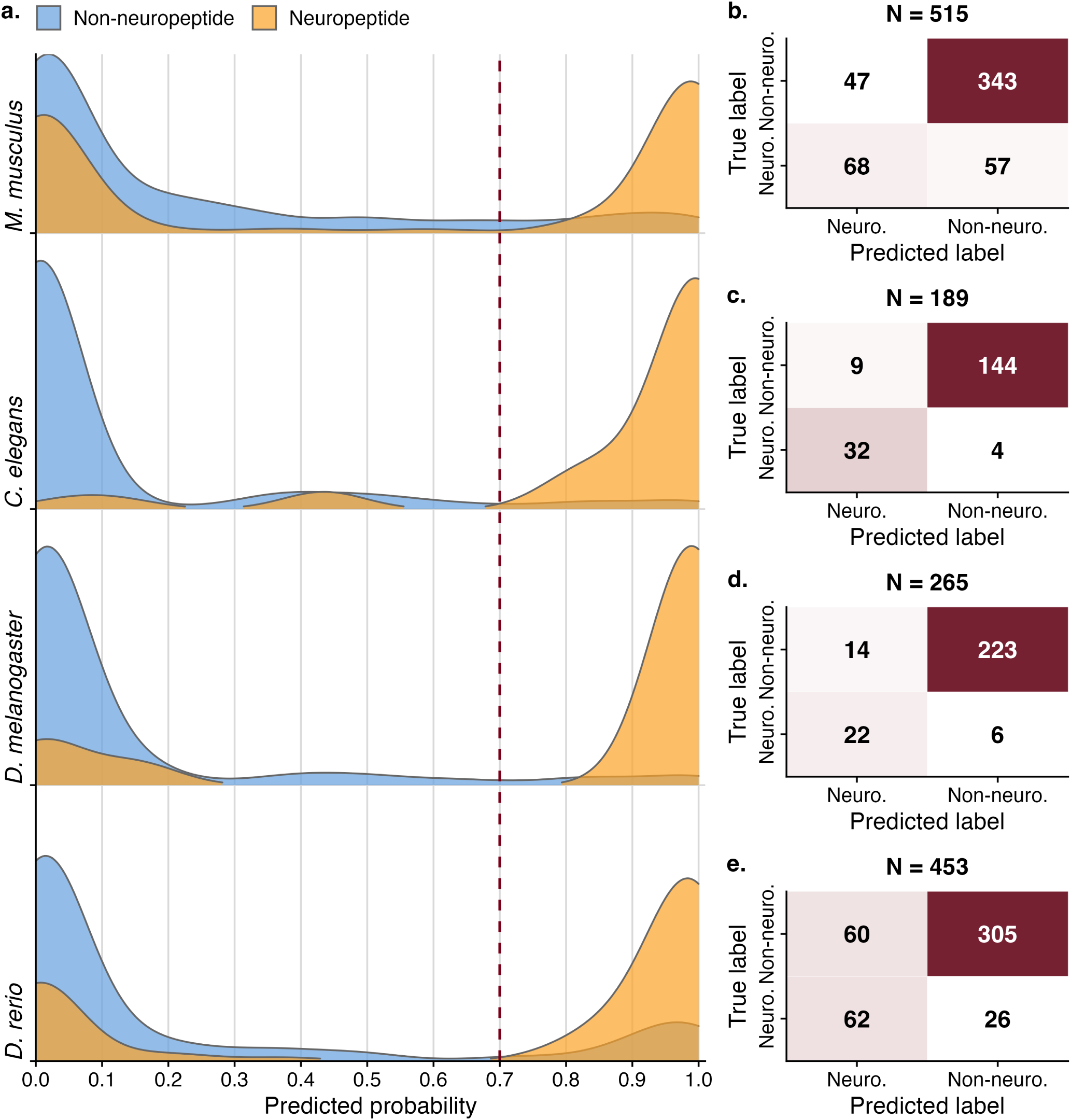
Generalization of InterPLM-NP to four additional secretomes. (a) Ridgeline density plots of predicted neuropeptide probabilities for annotated neuropeptides and non-neuropeptide secreted proteins across four species (*M. musculus*, *C. elegans*, *D. melanogaster*, and *D. rerio*). The red dashed line indicates the probability threshold of 0.7 used to binarize predictions for confusion-matrix calculations in panels b–e. (b) Confusion matrix for *M. musculus*. (c) Confusion matrix for *C. elegans*. (d) Confusion matrix for *D. melanogaster*. (e) Confusion matrix for *D. rerio*

In contrast, the *M. musculus* and *D. rerio* secretomes showed a bimodal neuropeptide probability distribution (Fig. 6a), with one group of annotated neuropeptides receiving low scores and another scoring highly. This pattern is consistent with greater heterogeneity in neuropeptide classes and annotation differences, which may cause some mouse and zebrafish neuropeptides to be less well captured by the features learned from the training set. Nonetheless, in both secretomes the model shows strong discrimination for a large subset of neuropeptides. By comparison, the fly and worm secretomes contain fewer, less diverse neuropeptides, and their probability distributions are less complex.

Confusion-matrix analyses further quantified performance across species (Fig. 6b–e). For *C. elegans*, InterPLM-NP achieved high recovery of annotated neuropeptides (TP=32, FN=4) with few false positives (FP=9; TN=144; Fig. 6c). Similarly, performance in *D. melanogaster* was also strong (TP=22, FN=6; FP=14; TN=223; Fig. 6d). In *M. musculus*, the model recovered many neuropeptides (TP=68) but also missed a substantial number (FN=57), largely consisting of larger protein hormones such as growth hormone and prolactin, and produced a moderate number of false positives (FP=47; TN=343; Fig. 6b), matching the bimodal neuropeptide distribution observed in the ridgeline plot. The same pattern was observed in *D. rerio* (TP=62, FN=26; FP=60; TN=305). These confusion-matrix counts are also summarized by accuracy, precision, recall, and *F*_1_ scores at the 0.7 probability threshold across all four species (Table 3).

**Table 3:**
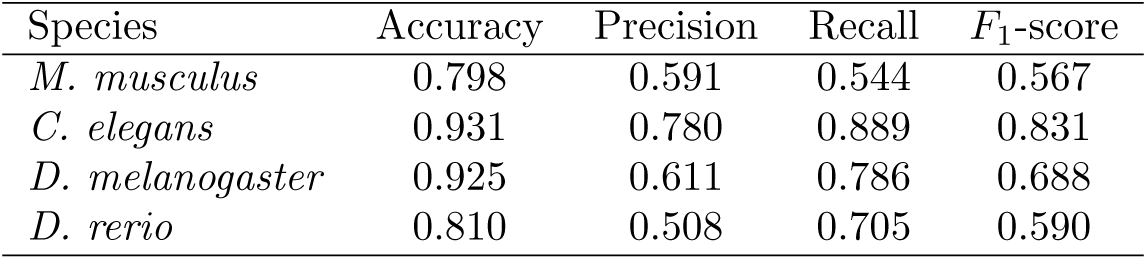
Performance of InterPLM-NP across four species’ secretomes at a probability threshold of 0.7.

| Species | Accuracy | Precision | Recall | $F_1$-score |
| --- | --- | --- | --- | --- |
| *M. musculus* | 0.798 | 0.591 | 0.544 | 0.567 |
| *C. elegans* | 0.931 | 0.780 | 0.889 | 0.831 |
| *D. melanogaster* | 0.925 | 0.611 | 0.786 | 0.688 |
| *D. rerio* | 0.810 | 0.508 | 0.705 | 0.590 |

Finally, we manually reviewed all secreted proteins that were predicted as neuropeptides (probability *≥* 0.7) but not originally annotated as neuropeptides across the four additional species. This review confirmed 19 (mouse), 8 (*C. elegans*), 8 (*D. melanogaster*), and 38 (*D. rerio*) of these candidates as neuropeptides missing from the original annotation, which were reclassified as true positives; the remaining candidates were confirmed as genuine false positives and retained their original label. Because only the entries predicted as neuropeptides were reviewed, this reannotation only moved entries from false positive to true positive and did not affect the false negative or true negative counts. Confusion matrices reflecting this reannotation are shown in Supplementary Fig. S4.

Together, these results show that InterPLM-NP generalizes well across species, with particularly strong performance in worm and fly secretomes and more bimodal behavior in mouse and zebrafish. It is important to note that these non-human secretomes were not explicitly held out or homology-filtered during training. Only the human secretome was held-out, so cross-species results should be interpreted as an additional generalization check, rather than an official benchmark.

## 4 Discussion

In this study, we built a pipeline to predict neuropeptides using sparse autoencoder (SAE) features from InterPLM. By converting dense ESM-2 embeddings into sparse features, we were able to identify individual features that were strongly associated with neuropeptide labels and use them to train a logistic regression classifier. Overall, top InterPLM (SAE) features captured strong neuropeptide-related signal, and neuropeptides tended to cluster toward an extreme area of the InterPLM embedding space within the human secretome. Using the top-ranked features, we trained a logistic regression model (InterPLM-NP) and selected a final top feature set of 320 features that correlate with neuropeptides. The resulting classifier performed well on the curated human secretome, where high-probability predictions were strongly enriched for annotated neuropeptides and additionally outperformed two existing neuropeptide prediction tools. When applied to additional secretomes, InterPLM-NP also generalizes across species, with strong performance in *C. elegans* and *D. melanogaster* and more mixed results in *M. musculus* and *D. rerio*. Together, these findings show that InterPLM SAE features provide a practical way to leverage protein language model embeddings for neuropeptide prediction.

Computational peptide discovery has traditionally relied on sequence homology searches (e.g., BLAST) and hand-crafted rules such as motif scans, signal peptide detection, and cleavage-site detection [2, 33, 35]. While often effective, these approaches can struggle when peptides are short, fast-evolving, or lack conserved sequence features, which is common for many bioactive peptides and neuropeptides [20, 6]. More recently, protein large language models and other embedding-based approaches have offered an alternative by learning sequence representations that capture biochemical and functional similarity beyond alignment-based homology [26, 19, 16, 4, 22, 17, 38, 39], enabling embedding-based search and annotation in diverse peptide-discovery contexts, [40], [14], [12]. In parallel, multiple protein or peptide predictors now incorporate embeddings from PLMs to improve prediction accuracy [9, 40, 13, 14, 43, 4]. A key remaining challenge is interpretability. Dense embeddings often encode many overlapping concepts, making it difficult to connect individual features to biological meaning [4, 1]. InterPLM addresses this by training sparse autoencoders (SAEs) on ESM-2 embeddings to produce sparse, more interpretable features, and subsequent work has explored how latent features can support functional annotation [29, 28, 1]. Our results build on this line of work by showing that SAE-derived features can be directly ranked for neuropeptide association and then combined in a simple classifier, providing a practical bridge between dense protein language model embeddings and feature-level interpretability [29].

Another advantage of sparse feature representations is their potential to improve learning in low-data (small-N) settings, where assembling large, well-labeled biological datasets is difficult. Recent work has shown that sparse autoencoder features can be particularly effective for low-N protein function prediction because they compress information from pretrained embeddings into a smaller set of task-relevant features that can be learned from far fewer labeled examples [37]. In practice, focusing a classifier on a compact set of informative features can reduce training time and the amount of labeled data needed to achieve useful performance. This is directly relevant for neuropeptide discovery, where high-confidence annotations for neuropeptides remain limited and uneven across species. In this context, SAE-based feature ranking paired with simple classifiers offers a promising pipeline to extract predictive signal from protein language model embeddings and prioritize candidates even when only modest labeled datasets are available.

Future work can expand this pipeline from a neuropeptide predictor into a more complete secretome annotation tool. One possible extension is to apply the same SAE-based feature ranking and classifier training framework to cleavage-site prediction by learning which sparse features best distinguish cleavage-associated residues and then training residue-level models on those features. Because high-quality cleavage annotations are relatively scarce (small-N), this task is well suited to approaches that concentrate signal into a smaller and more learnable set of features. In parallel, extending the model to predict post-translational modifications (PTMs) that are common in neuropeptides such as C-terminal amidation would also further improve potential neuropeptide candidate prioritization [41]. Downstream work could also focus on subclassifying predicted precursors into neuropeptide families or functional categories including receptor-target classes. Collectively, the features learned for these tasks could be integrated into a modular annotator that outputs multiple labels (e.g., neuropeptide, predicted cleavage sites and mature peptides, PTMs, and family-level classes), enabling more robust biological interpretation and more promising experimental follow-up.

## Supplementary Figures

**Figure S1:**
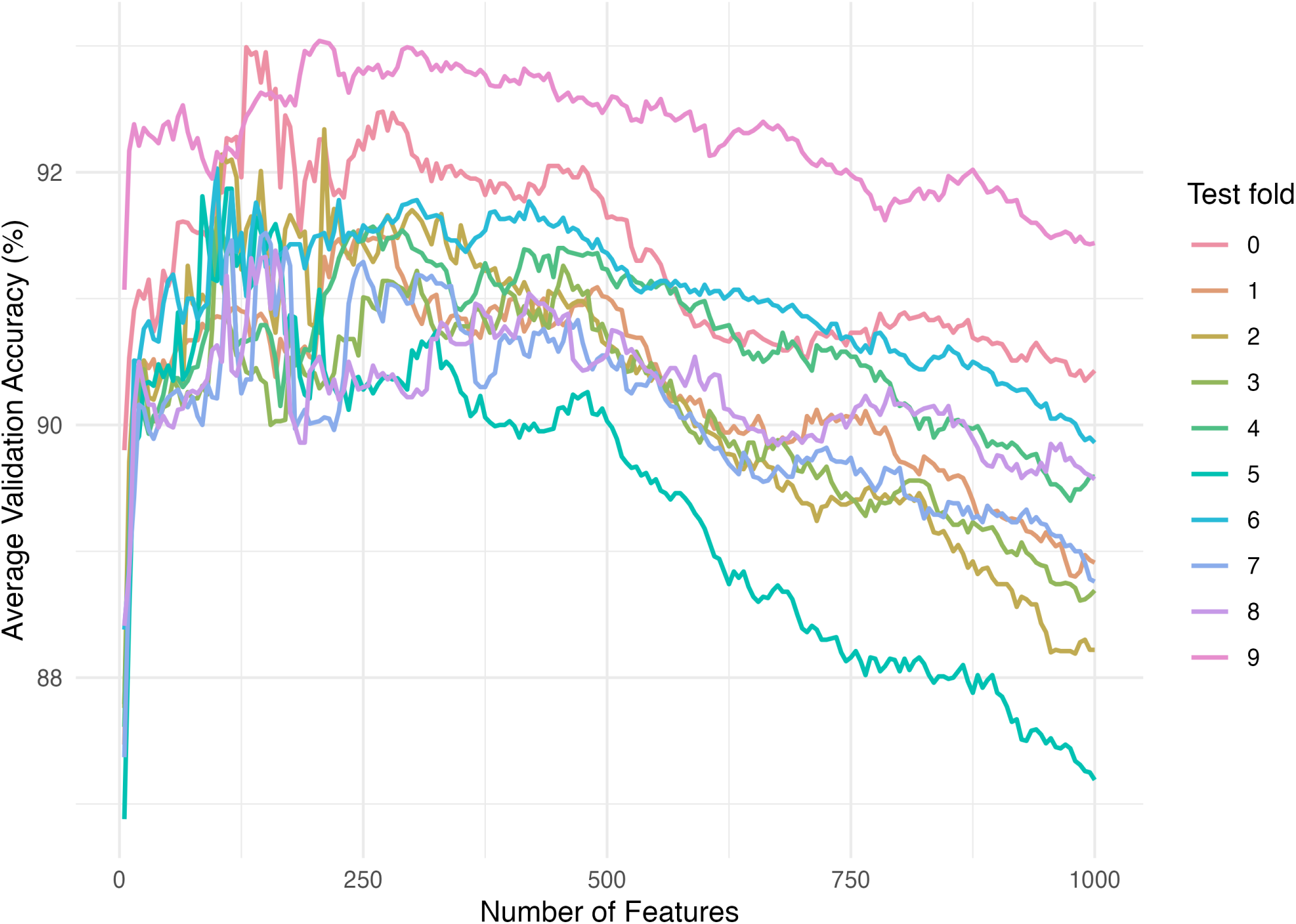
Average validation fold accuracy across feature set sizes. In a 10-fold cross-validation, each fold was consecutively held out as the test fold. For each held-out fold, the remaining 9 folds were used for validation, and their accuracies were averaged for each number of features *n*, evaluated in increments of *n* = 5 features up to a maximum of *n* = 1000. Colors indicate which fold was excluded from averaging and reserved for subsequent test accuracy evaluation.

**Figure S2:**
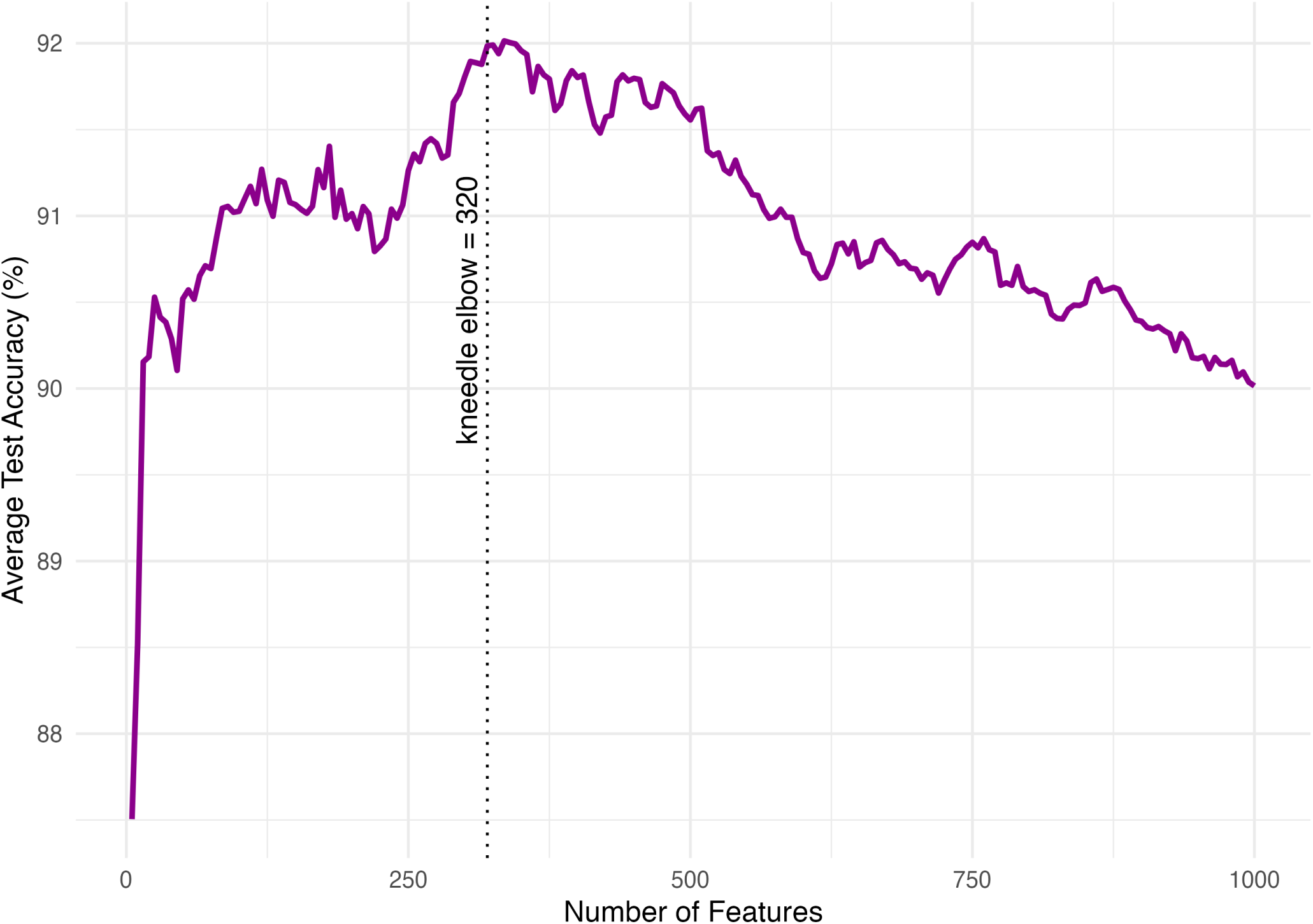
Average test fold accuracy across feature set sizes. In a 10-fold cross-validation, each fold was consecutively held out as the test fold, and its accuracy was evaluated using the model trained on the remaining 9 folds. Accuracies were averaged across all test folds for each number of features *n*, evaluated in increments of *n* = 5. The optimal number of features was found using the Kneedle algorithm [27].

**Figure S3:**
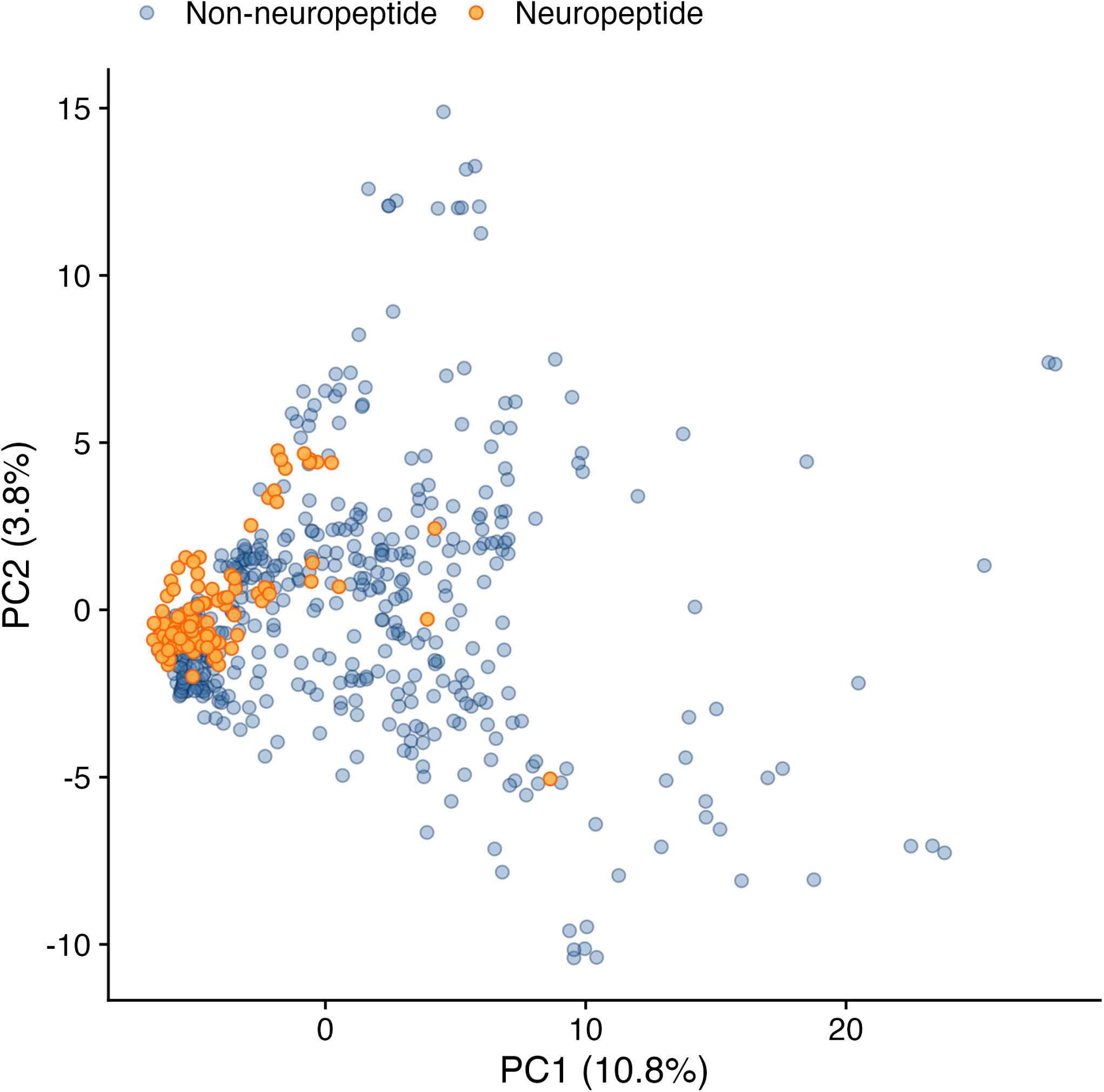
PCA of human secretome proteins in the space of the top 320 InterPLM (SAE) features associated with neuropeptides. Orange points indicate annotated neuropeptides and blue points indicate non-neuropeptides.

**Figure S4:**
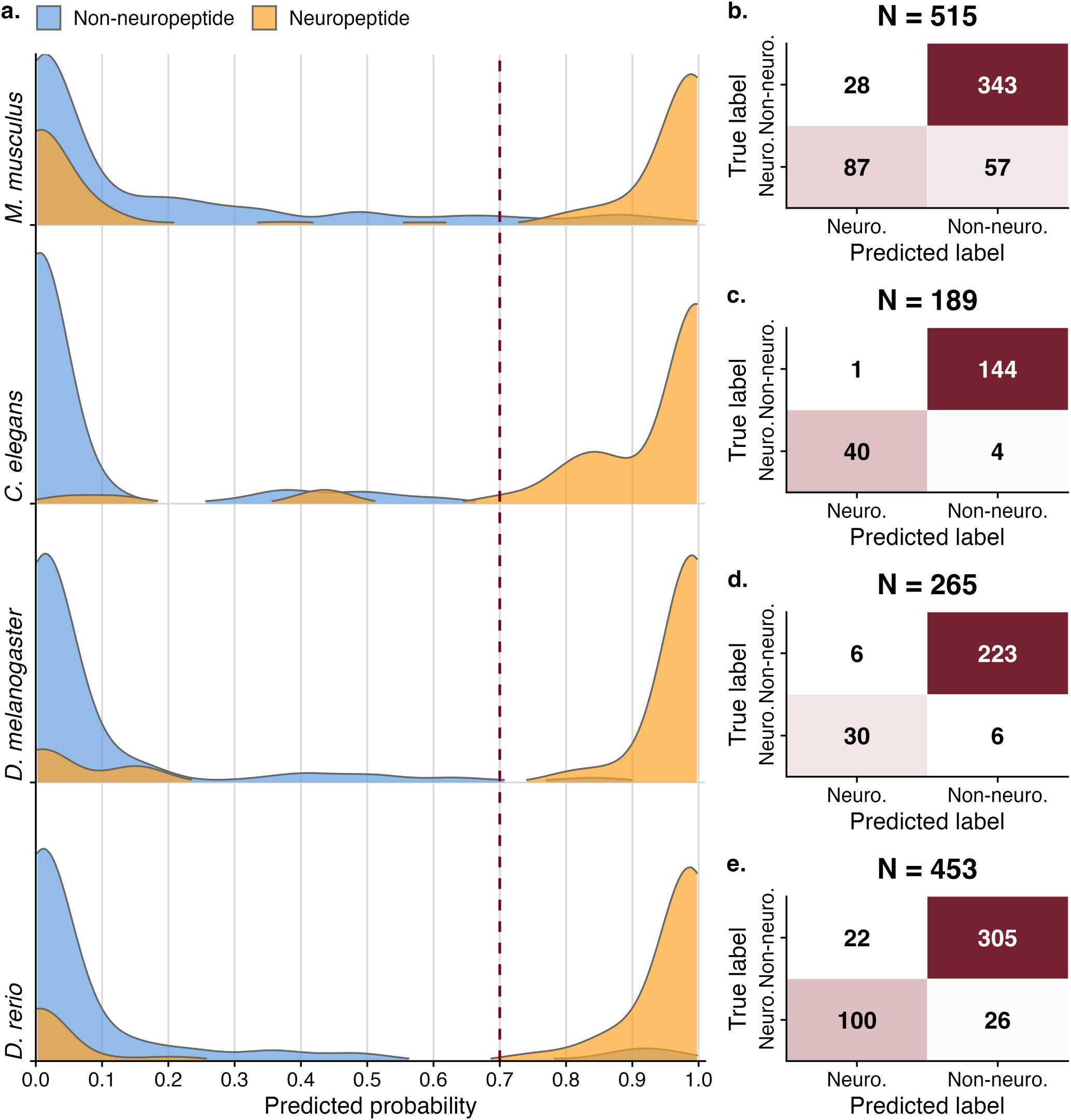
Performance of InterPLM-NP on four reannotated secretomes. (a) Ridgeline density plots of predicted neuropeptide probabilities for annotated neuropeptides and non-neuropeptide secreted proteins across four species (*M. musculus*, *C. elegans*, *D. melanogaster*, and *D. rerio*). The red dashed line indicates the probability threshold of 0.7 used to binarize predictions for confusion-matrix calculations in panels b–e. (b) Confusion matrix for *M. musculus*. (c) Confusion matrix for *C. elegans*. (d) Confusion matrix for *D. melanogaster*. (e) Confusion matrix for *D. rerio*

**Figure S5:**
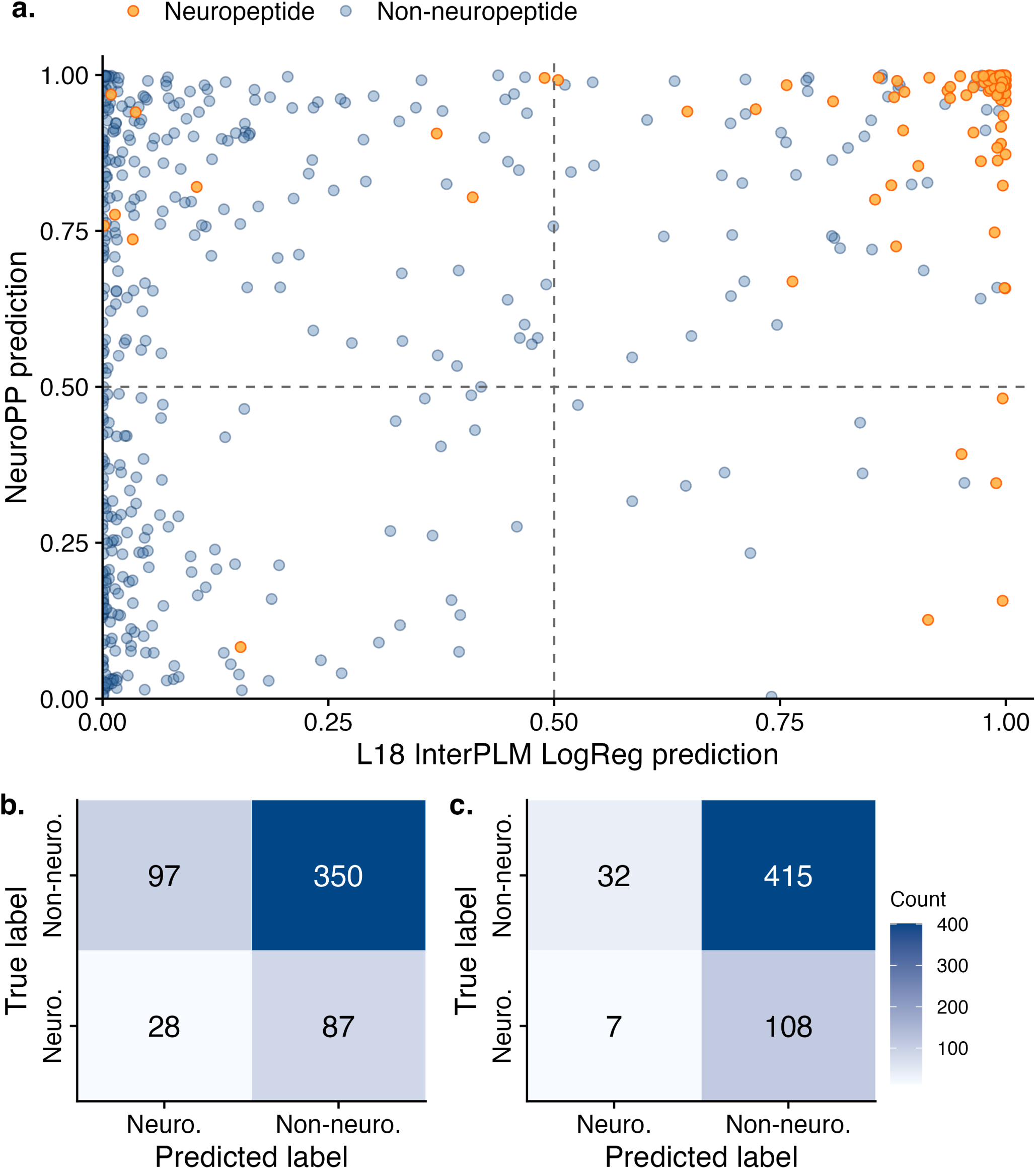
Performance of two additional baseline neuropeptide prediction models on the human secretome. (a) Correlation between InterPLM-NP probabilities (x-axis) and NeuroPP probabilities (y-axis), with Pearson correlation *r* = 0.447 (*p* = 6.96 *×* 10*^−^*^29^). (b) Confusion matrix for NeuroPred-FRL predictions (binary output). (c) Confusion matrix for NeuroPredPLM predictions (binary output). Note: models (b) and (c) were trained to predict mature neuropeptides rather than neuropeptide precursor peptides.

**Figure S6:**
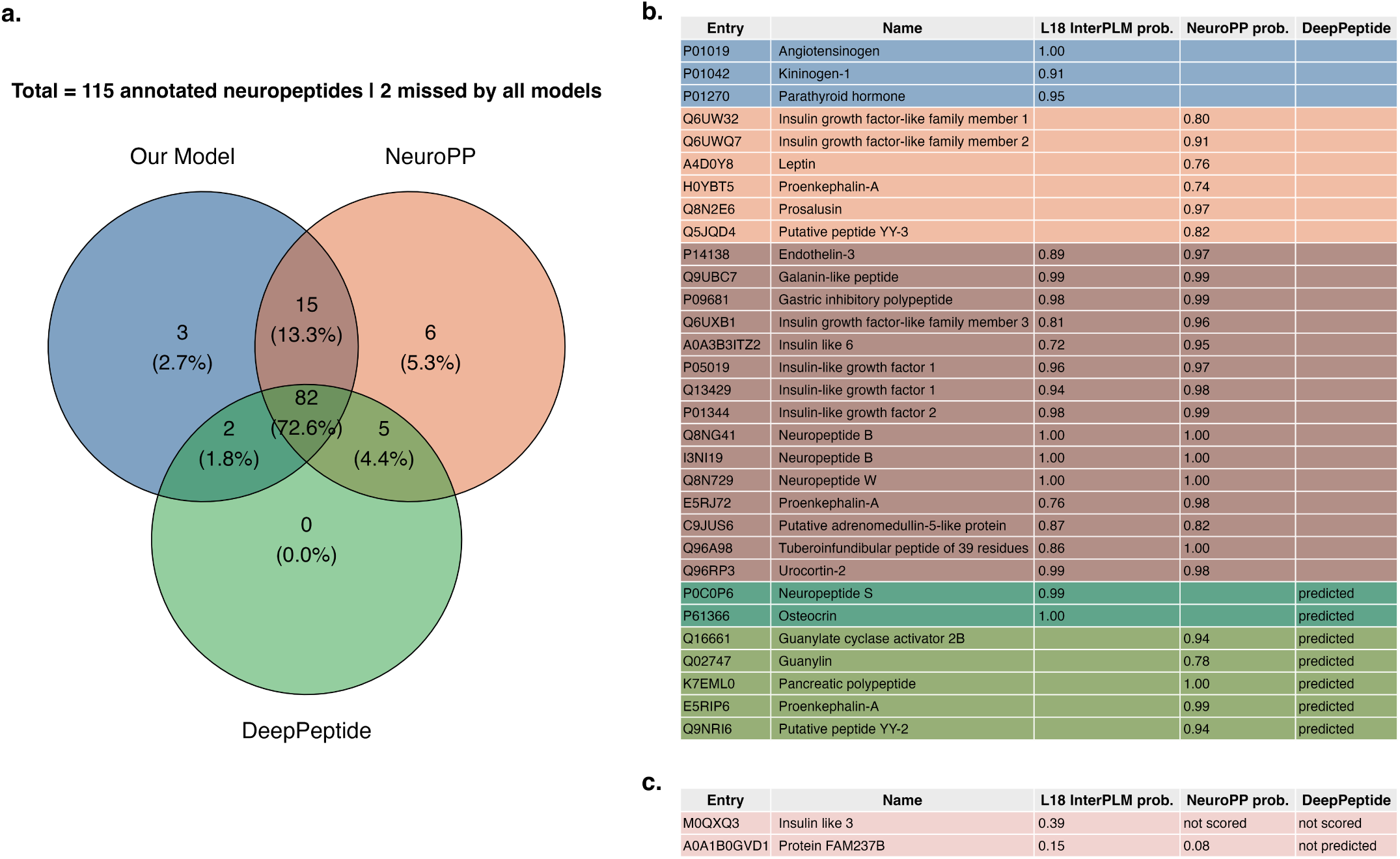
Comparison of neuropeptide predictions across models on the annotated human secretome. (a) Venn diagram showing how many annotated human secretome neuropeptides were correctly predicted by each model. Overlaps indicate agreement between models. (b) Predicted neuropeptides for each partial-overlap region of the Venn diagram, with each model’s predicted probability; entries and probabilities are colored by which combination of models predicted them. The 82 entries found by all three models are omitted here for space and are shown only as a count in (a). (c) The two annotated neuropeptides not predicted by any of the three models.

## Supplementary Data

- Supplementary Data File S1: Human secretome predictions (`human predictions.csv`)
- Supplementary Data File S2: Mouse secretome predictions (`mouse predictions.csv`)
- Supplementary Data File S3: Drosophila melanogaster secretome predictions (`fly predictions.csv`)
- Supplementary Data File S4: Caenorhabditis elegans secretome predictions (`worm predictions.csv`)
- Supplementary Data File S5: Danio rerio secretome predictions (`zebrafish predictions.csv`)

